# Estimation of Vertex-wise Sulcal Width Maps on Cortical Surfaces

**DOI:** 10.1101/2022.11.09.515775

**Authors:** Alberto Fernández-Pena, Francisco J. Navas-Sánchez, Daniel Martín de Blas, Luis Marcos-Vidal, Pedro M. Gordaliza, Isabel Martínez-Tejada, Joost Janssen, Susanna Carmona, Manuel Desco, Yasser Alemán-Gómez

## Abstract

Sulcal width, defined as the physical distance between opposing sulcal banks, has shown promise as a biomarker. We present the first method to obtain a vertex-wise representation of this metric directly on the brain’s cortical surface. The algorithm samples the surface at different depths and estimates the distances between the opposing sulcal banks. The method is validated against a simulated sulcus and compared to other sulcal width tools based on regions of interest, showing solid correlations between the proposed algorithm and two widely-used reference methods. A vertex-wise assessment of the sulcal width is carried out by evaluating the correlation between sulcal width and age in a sample of an aging population, revealing clusters in the central, cingulate, and temporal sulcus regions associating wider sulci for older participants. The results confirm that the algorithm is reliable for obtaining sulcal width maps, allowing for vertex-wise analyses, and providing aggregated measures similar to the existing methods. The present algorithm is publicly available via https://github.com/HGGM-LIM/SWiM.

## Introduction

The sulcal morphology of the human brain cortex can be quantified using magnetic resonance imaging (MRI). Sulcal width, defined as the physical distance between opposing sulcal banks, has shown promise as a biomarker for brain changes in disease, maturation, aging, or pregnancy (1–9). Most current methods for assessing sulcal width are limited to the region of interest (ROI) level. The first automatic 3D ROI-based approach for sulcal width calculation was developed by Mangin et al. and implemented in the BrainVISA image processing suite (11). This method does not directly measure the physical distance between the sulcal banks. The method relies on the generation of a fold-specific medial mesh that follows the sulcal fundus and rises from the fundus to the intersection with a convex brain hull. The average sulcal width for a given fold is then approximated by the quotient of the volume of the cerebrospinal fluid (CSF) buried inside the hull-covered fold with the surface area of the medial mesh (12).

Another approach is presented by Kochunov et al., which also relies on the extraction of sulcal medial surfaces obtained from BrainVISA. The algorithm traces two opposite vectors along the normal direction up to their intersections with the sulcal bank for each point on the sulcal medial mesh. These vectors intersect on both sides of the sulcal basin, and the Euclidean distance between both intersection points is considered the sulcal width for a given medial surface point. All sulcal width values are averaged to obtain one mean sulcal width measure for each fold. While this method is a more direct measure of the distance between sulcal banks than the one presented in Mangin et al., it has the same limitation as the previous algorithm: it only provides an average width value for a given fold (ROI).

Similarly, Madan proposes a method to obtain a sulcal width measure based on the standard outputs obtained by FreeSurfer (15, 16) while avoiding the calculus of a medial surface of the folds. Here, opposing vertices in the boundaries between sulcal and gyral regions in the Destrieux cortical parcellation (17) are detected. The final metric is the median of the Euclidean distances between these opposite vertices, providing a single value for each ROI. This method depends on the gyri/sulci definition proposed by the Destrieux atlas. The sulcal width measures are only obtained in the gyri/sulci boundary, relying on the accuracy of the mapping of the sulcal labels to the individual cortical anatomy.

A voxel-based sulcal width estimation approach is presented in Mateos et al.. The algorithm uses a voxel representation of the pial surface and estimates voxels located in the center of the sulcal space using the Euclidean Distance Transform (EDT) (19) and a Local Maxima detection algorithm. The sulcal width is given by double the distance to those estimated voxels. This method is limited by the possible discretization artifacts and the resolution selected for this discretization. Also, in their provided analyses, the method does not supply a value for each voxel in the pial boundary, but rather an aggregated metric for each region, remaining unclear how the values can be projected to the cortical surface. Additionally, the lack of an accessible implementation of the method hampers its usage.

Complementing prior methods, a vertex-wise assessment of sulcal width would enable quantification of the effects of width and its relationship with other morphological and non-morphological data at high spatial resolution. Furthermore, vertex-wise estimates of sulcal width would also allow for the mapping of width along the sulcal banks’ depth to the sulcal fundus. Normalization of such profile maps may provide new clues about, e.g., at which sulcal depth sulcal widening is more pronounced and whether diseases are associated with changes in profile maps (20).

In the present work, we present a novel and robust method that provides a vertex-wise sulcal width map estimation based on the estimation of isolines at different depth levels. Then, the points in each isoline are paired with the corresponding on the opposite sulcal wall. The distances between these points are used to estimate the vertex-wise sulcal width map. The proposed approach tackles the main issues from prior methods: 1) contrary to prior methods (10, 13), it does not rely on a medial surface of the sulci to obtain the sulcal width map; 2) the method produces full vertex-wise maps for the sulcal width instead of sulcal averages (10, 13, 14); 3) the estimation computes the sulcal width by sampling the surface at different depth levels; 4) the method works entirely over the pial mesh triangulation, avoiding possible artifacts derived from the voxelization of the cortical surface (18), and 5) the method is entirely compatible with standard FreeSurfer output.

The performance of the developed algorithm is assessed in two different ways: First, using a synthetic (computed generated) sulcus surface as a ground truth. Second, comparing the reproducibility and distribution of averaged sulcal width values obtained using MRI data from the Human Connectome Project (HCP) (21) between the proposed algorithm and two well-known, publicly available methods: BrainVISA (10) and Kochunov et al.. Finally, a relationship between aging and the increase of average ROI-based sulcal width has been described (6, 22–24). However, it is not clear whether the age association is constant across sulci or whether there are focal regions that show a stronger association. We, therefore, present a vertex-wise analysis of the correlation between sulcal width and age in a subset of the Open Access Series of Imaging Studies (OASIS) (25).

The method is freely available in (https://github.com/HGGM-LIM/SWiM).

## Methods

### Cortical Surface Extraction and Sulcal Depth Computation

The algorithm begins with the surface extraction process from T1w anatomical images to generate a triangular mesh that wraps the boundary between grey and CSF tissues, i.e., the pial surface. The obtained mesh comprises vertices connected by edges where a set of three closed edges defines a face. In this work, this process is performed by the FreeSurfer pipeline (26, 27). However, this methodology can be applied to triangular meshes generated using any other image processing software such as BrainVISA (11) or BrainSuite (28).

Once the cortical mesh is obtained, the first step is to estimate a vertex-wise depth map of the pial surface. In this case, we are using travel depth, understood as the shortest distance from any vertex to the convex hull without intersecting the surface (29). The travel depth is computed using Mindboggle^1^ (30). Travel depth is selected over geodesic depth because the latter tends to overestimate the metric, especially in areas such as the insula (30).

### Depth-based Isolines Construction and Labeling

The designed algorithm samples the local characteristics of the generated pial surface at different depth levels. For this, a set of isolines is created using the mentioned surface and its corresponding depth map^2^.

These isolines are a collection of closed curves over the individual surface, computed for a discrete set of equispaced depth levels starting from a minimum initial depth (Figure 1). The distance between depth levels should be chosen to obtain a set of isoline points denser than the surface vertices, guaranteeing that every edge on the surface is thoroughly covered but without increasing the computational costs unreasonably (see S1 of the Supplementary Material). A reasonable value for this parameter is 1/5 of the average edge length of the surface (approximately 1 mm for the pial surfaces obtained from FreeSurfer). Therefore, for this work, we create a depth profile of sulcal depth values for every 0.20 mm. We also set the initial depth value to generate the isolines to 1.5 mm to avoid crests of the gyral regions where the definition of sulcal width is unclear.

**Fig. 1.**
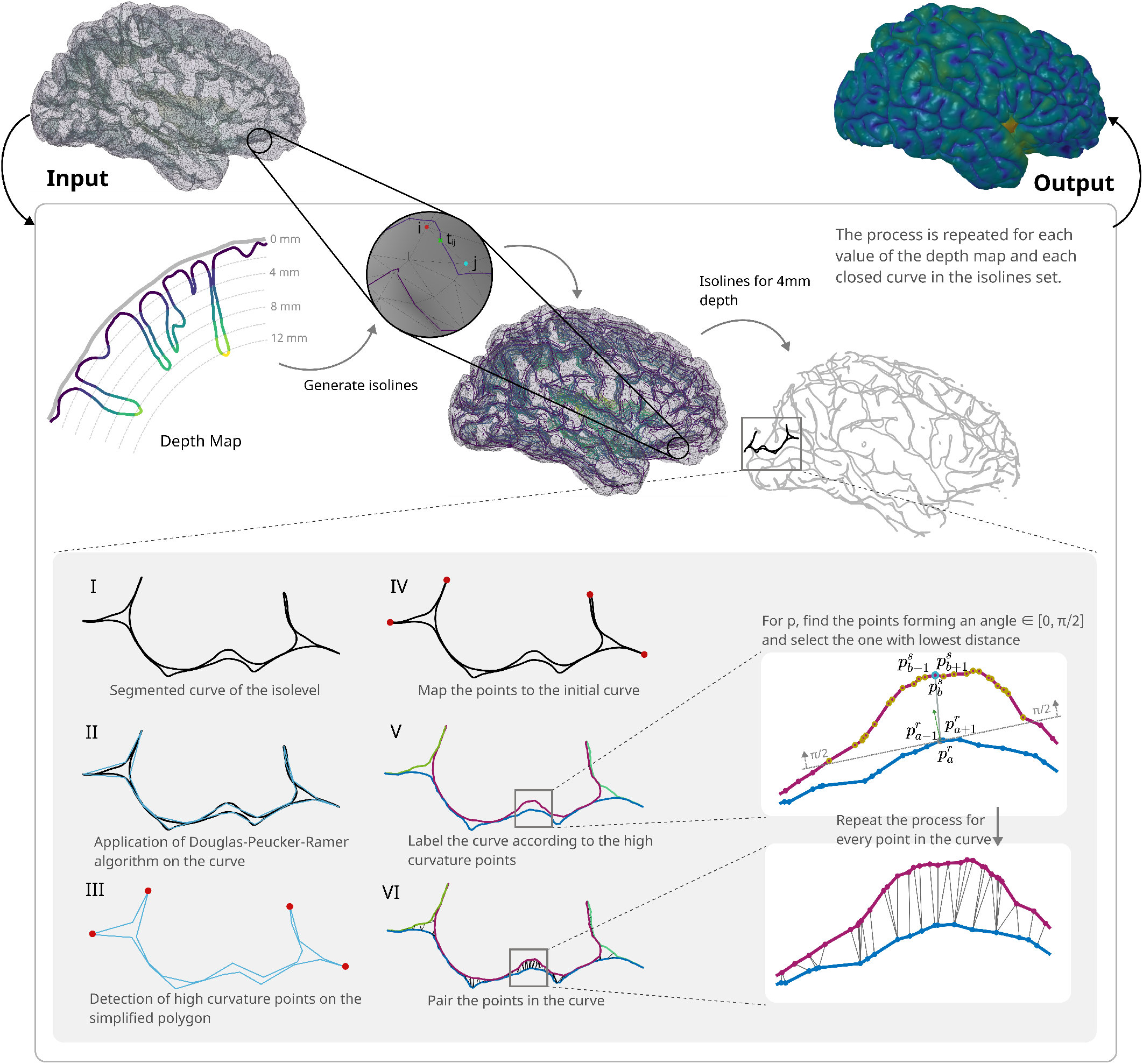
Graphic representation of the method steps. The algorithm takes the pial surface as input, generates the depth map, and for each depth level, the resulting isolines are processed as represented in the gray box. Finally, the output is a vertex-wise sulcal width map generated by aggregating all the width measures for each vertex.

For each depth level, we obtain the points where the isoline crosses the surface edges (see the zoomed input map in Figure 1) by computing the following ratio for each surface edge:

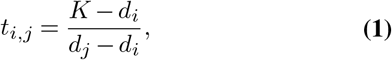

where *K* is the depth value for the isoline, *i* and *j* are any pair of vertices connected by the edge, and *d*_*i*_ and *d*_*j*_ are the corresponding depth values. The resulting *t*_*i,j*_ is the parameter where the depth value *K* falls over the line defined by the edge with vertices *i* and *j*. Only values of *t*_*i,j*_ ∈ [0, 1] represent valid intersections between the edges and the isoline; hence, the edges yielding values out of that interval are discarded. Finally, the positions in which the isoline intersects the edges are computed as *p*_*i,j*_ = *v*_*i*_ + *t*_*i,j*_(*v*_*j*_ *− v*_*i*_), where *v*_*I*_ denotes the position of the *i* vertex.

Then, each closed curve in the isoline is segmented attending to the sulcal banks where it lies. To do this, we need to identify the points where the change of sulcal wall occurs, which would be those points with a high curvature value. As the curves are continuous and contain a high number of points (Figure 1-I), the angles between consecutive points tend to be very plain, making it challenging to identify sharp angles. This problem is tackled using a line simplification algorithm to transform the curve into a polygon (31, 32) (Figure 1-II). This algorithm decimates a curve by selecting a minimum number of points from a given shape based on the maximum distance between the original curve and the simplified polygon. Supposing the new polygon is formed by the points {*r*_1_, *r*_2_, …, *r*_*n*−1_, *r*_*n*_ }, the cosines of the angles between the consecutive segments in the resulting simplified polygon are estimated using the dot product between those corresponding vectors 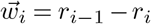 and 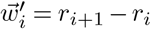.

The polygon points forming an angle lower than 3/5*π* with their neighbors are considered the points that mark the sulcal bank change, as wider angles are considered an effect of the normal sulcal sinuosity (Figure 1-III). This parameter was selected in a heuristic manner after a visual inspection of different subjects. Then, as there is correspondence between the simplified polygon and the original curve, these high curvature points are marked on the initial curve (Figure 1-IV). As these high curvature points divide the curves according to their sulcal banks, the algorithm assigns a different label to each curve section enclosed between two markers (Figure 1-V). Those labels are used in the next step to match points in different labels and estimate the sulcal width.

The label segmentation process is repeated for each closed curve of each depth-level isoline.

### Sulcal Width Computation

The isoline *I* is composed of a set of *N* points, 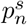, organized in *S* labels, where *s* ∈ {1,…, *S*} denotes the label to which the point belongs, and *n* ∈ {1,…, *N* } represents the index of the point in the isoline. We pair each isoline point, 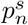, with its corresponding in the opposite sulcal bank, *q*_*n*_. For this, we first define the set of candidate points, 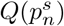 as:

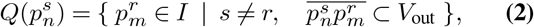

where 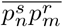 denotes the line segment, connecting the points 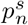 and 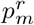, and *V*_out_ the volume outside the pial surface. To obtain the set of candidates, 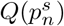, first, we restrict the candidate points to those on a different label where the angle between the normal of the surface at 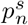 and the vector 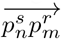 is in the range of [0, *π/*2] (see yellow points in Figure 1-V close-up). This ensures that the segment 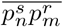 goes outwards the surface, as otherwise it would measure gyral span. Then, the intersections between the segment 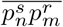 and the pial sursurface are calculated (33). The connections where 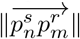 is greater than the distance from 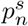 to the first surface intersection in the direction of 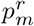 are discarded, as the segment is intersecting and crossing the pial mesh. From these candidate points, we select the corresponding point, *q*_*n*_, on the opposite sulcal bank as the one with minimum Euclidean distance, i.e.,

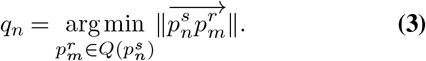

Repeating this process for every point in the isolines generates a list of point pairs where the Euclidean distance among each pair of points is computed and imputed to the vertices in the pial surface to obtain the sulcal width map. As each point in the isolines lies in a mesh edge, we assign the distance 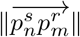 to each of the vertices forming that mesh edge. Most of the vertices in the surface have contributions from multiple isolines, so the final width value for each vertex takes the median value of all the contributed width measures. The isolines are generated using a depth step small enough to have multiple isoline values contributing to each vertex; hence, most vertices in the width map should be covered. For those that remain unlabeled (primarily those in the upper gyral regions), the values are interpolated by iteratively averaging the values of their neighbors.

Finally, we apply a linear smoothing step to the width map to refine the possible discontinuities that can appear in some regions. This smoothing averages the value of each vertex with those in its neighbors to ensure spatial coherence.

### Validation

#### Validation Using Simulated Data

A triangular mesh mimicking a cortical sulcus is created to assess the performance of the proposed method. Parameters such as sulcal length, width, or depth profile are used to reproduce the morphology of a cortical sulcus, serving as a control item for the quality and reliability of the vertex-wise maps.

This synthetic sulcus has a total length of 25 cm with a spatial resolution of 10 points/cm along the longitudinal axis. For each point of the longitudinal axis, a Gaussian distribution is generated and placed orthogonally to this axis with its mean spatially located over the point (see Figure 2A). The width and depth of the simulated surface along the longitudinal axis are adjusted by altering the parameters of these Gaussian distributions. First, the standard deviation (*σ*) is varied to modify the width across its longitudinal axis. This variation is generated by a cosine function between 0 and 2*π*. Finally, depth fluctuation is set by multiplying the Gaussian distribution by a different amplitude factor at each longitudinal point. Both width and depth variations throughout *σ* and the amplitude parameters create a narrow and shallow sulcus in the sharp ends that rapidly gain width and depth and then lightly decreases again in the central region. Figure 2B depicts the resulting shape of the simulated sulcus

**Fig. 2.**
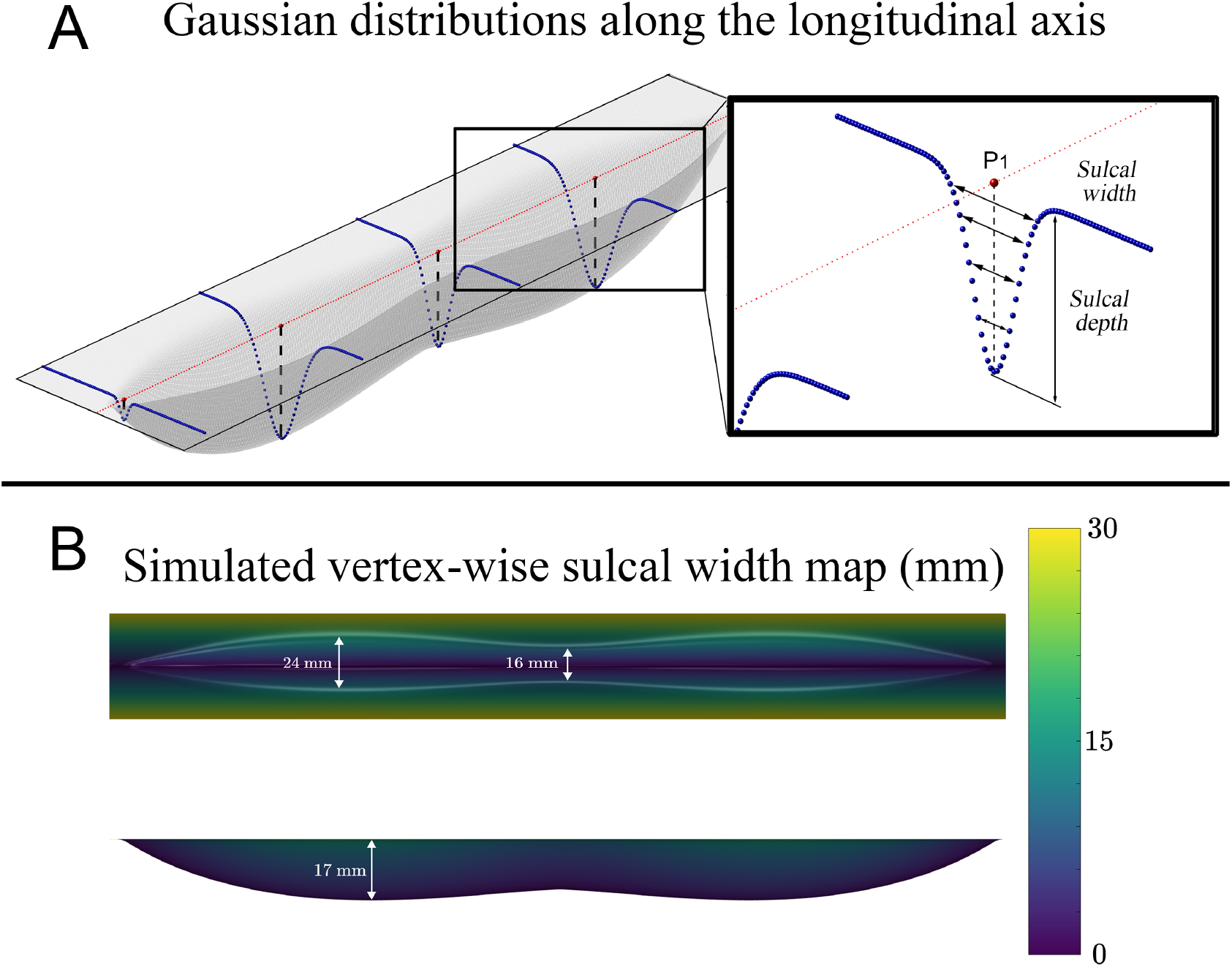
**A:** Schematic view of the generation of the simulated sulcus. **B:** Top and front perspective views of the simulated sulcus. The color bar and color map show the simulated width values in millimeters. Annotations show the main dimensions of the sulcus.

As the Gaussian distributions are symmetric, pairs of points belonging to the same Gaussian distribution, located at the same depth but in opposite sulcal banks, are paired, and the Euclidean distance between them is taken as their reference (ground truth) sulcal width value (see Figure 2A). Finally, the mesh is generated by creating triangular faces linking each point to its corresponding top and left neighbors.

The resulting mesh is fed into the width estimation algorithm to create the vertex-wise width map. This map is compared to the reference sulcal width map. Only vertices with depth over 0.1 mm are used for the comparisons to avoid the top flat region that is not considered part of the sulcus.

#### Validation Using HCP Test-Retest Dataset

The proposed algorithm is also validated through comparison against two well-known sulcal width (or fold opening) estimation methods: BrainVISA (10) and Kochunov (13).

The data used for this validation is provided by the HCP Test-Retest dataset^3^ (21). This dataset includes 45 healthy subjects (13 male, age range of 22–35 years old) scanned twice (4.7 ± 2 months interval, minimum = 1 month, maximum = 11 months) on a Siemens 3T Skyra scanner in Washington University or University of Minnesota. T1w sagittal images were acquired using a Magnetization-Prepared Rapid Acquisition Gradient Echo (MPRAGE) sequence with 3D inversion recovery, echo time (TE) = 2.14 ms, repetition time (TR) = 2400 ms, inversion time (IT) = 1000 ms, flip angle (FA) = 8°, field of view (FOV) = 180 × 224 × 224 mm^3^, voxel size = 0.7 × 0.7 × 0.7 mm^3^. The HCP consortium also provides the cortical surfaces (pial and white) and their curvature maps, obtained by a project-specific FreeSurfer pipeline (34).

The pial surfaces are imported to the BrainVISA’s Morphologist (11) pipeline, where the medial sulcal surfaces are created. The medial sulcal surfaces are automatically labeled according to the sulcal region they occupy (35) and are manually corrected to minimize the individual differences in labeling between test and retest cortical surfaces. These medial meshes are projected over the gyral banks to generate sulcal basins corresponding to each labeled sulcus. This way, we generate corrected sulcal region maps for every subject.

All analyses are performed in ten primary sulci (five for each hemisphere) that are consistently present across individuals and show less inter-individual variability than other sulci: central sulcus, calcarine, superior temporal sulcus, parietooccipital fissure, and cingulate sulcus on both hemispheres (36).

For each of these sulci, we evaluate the performance of the proposed method for computing sulcal width values compared to the ones obtained by the other two methodologies (BrainVISA and Kochunov). These methods provide a unique sulcal width value for each sulcus. Hence, we average the width map generated by the proposed method in each vertex belonging to the same sulcal basins extracted from the Morphologist pipeline, discarding the points relative to gyral regions (i.e., vertices with depth below 1.5 mm).

#### Application to the OASIS-3 Dataset

To demonstrate the potential usages of the tool, the algorithm is applied to a cohort of adults drawn from the OASIS-3 dataset^4^ (25). The selected subset contains individual T1w MR images of 362 subjects, ages 42 to 95 years old. These images were acquired at the same Siemens TrioTim 3T scanner using a 3D GR/IR scanning sequence with the following parameters: echo time (TE) = 3.16 ms, repetition time (TR) = 2400 ms, inversion time (IT) = 1000 ms, flip angle (FA) = 8°, field of view (FOV) = 176 × 256 × 256 mm^3^, voxel size = 1 × 1 × 1 mm^3^.

This analysis aims to assess, vertex-wise, the effect of age over the sulcal width. FreeSurfer (v.7.2) (26, 27) is used to extract the cortical surfaces (white and pial) and to perform all statistical analyses. The individual sulcal width maps are estimated using the FreeSurfer outputs and then registered to the *fsaverage* surface. A geodesic Gaussian smoothing with a full width at half maximum (FWHM) of 5 mm is performed previous to the statistical analyses.

### Statistical Analyses

#### Simulated Data

Pearson’s correlation coefficient and a Bland-Altman plot are used to assess the estimated sulcal width map’s accuracy compared to the ground-truth values.

#### Test-Retest Dataset

The mean width values of each of the ten sulci were compared between different methods using paired Student’s t-test analyses.

The similarities between methods are assessed by linear regression. The width values of the proposed method are regressed against the two control algorithms, and Pearson’s correlation coefficients are calculated.

Finally, the reproducibility of each method is measured using both the Intraclass Correlation (ICC) and the Root Mean Square Error (RMSE) between the width values computed for test and retest acquisitions. ICC describes how similar are the width measures in the test and retest acquisitions compared to the total variation of the width measures across all acquisitions and all subjects, while RMSE gives a notion of the magnitude of the error between test and retest values.

For each analysis, resulting p-values are corrected for multiple comparisons (10 comparisons, 5 for each hemisphere) using False Discover Rate (FDR) (37), considering *q* ≤ 0.05 as significant. All statistical analyses are performed using Python packages Pingouin (38), and Scipy (39).

#### Subcohort of OASIS-3 Dataset

A general linear model including gender and age as a factor and continuous covariate respectively is designed, and 10,000 Montecarlo permutations are used to obtain a Cluster-Wise Probability (CWP) (40) using FreeSurfer. A p-value *<* 0.0001 as the Cluster-Forming Threshold (CFT) for multiple comparisons correction is selected.

## Results

### Results on the Simulated Sulcus

Figure 3 shows the results of the comparison between the simulated and estimated data. The correlation coefficient between vertex-wise simulated and estimated width values is 1 (Figure 3A). In the Bland-Altman plot (Figure 3B), the differences between simulation and estimation are reported on the vertical axis, while their mean is depicted on the horizontal axis. This plot shows higher discrepancies in regions with lower sulcal width which are located near the extremities and the fundus of the simulated sulcus (Figure 3C).

**Fig. 3.**
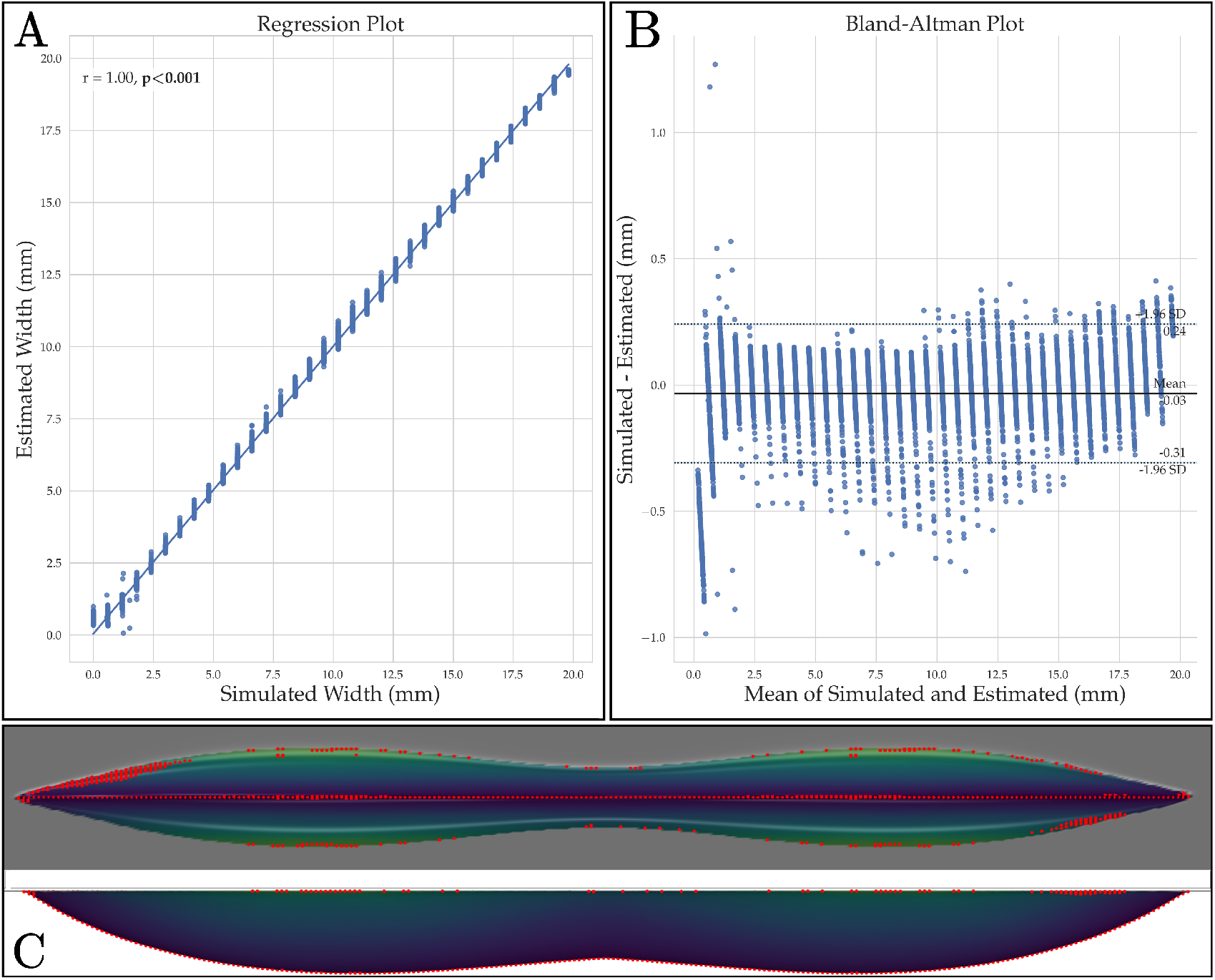
**A**: Regression plot between simulated and estimated width values of the vertices of the synthetic sulcus. **B**: Bland-Altman plot comparing simulated and estimated sulcal widths. **C**: Simulated sulcus with the points presenting width values out of the confidence interval highlighted in red. The gray region indicates depth values lower than 0.1 mm.

### Results on the HCP Test-Retest Dataset

For each of the ten sulci of interest, we can observe the mean width distribution in Figure 4 for the three inspected methods. Table 1 reports the pairwise comparison results of the distribution differences between the proposed method and the two control methods.

**Fig. 4.**
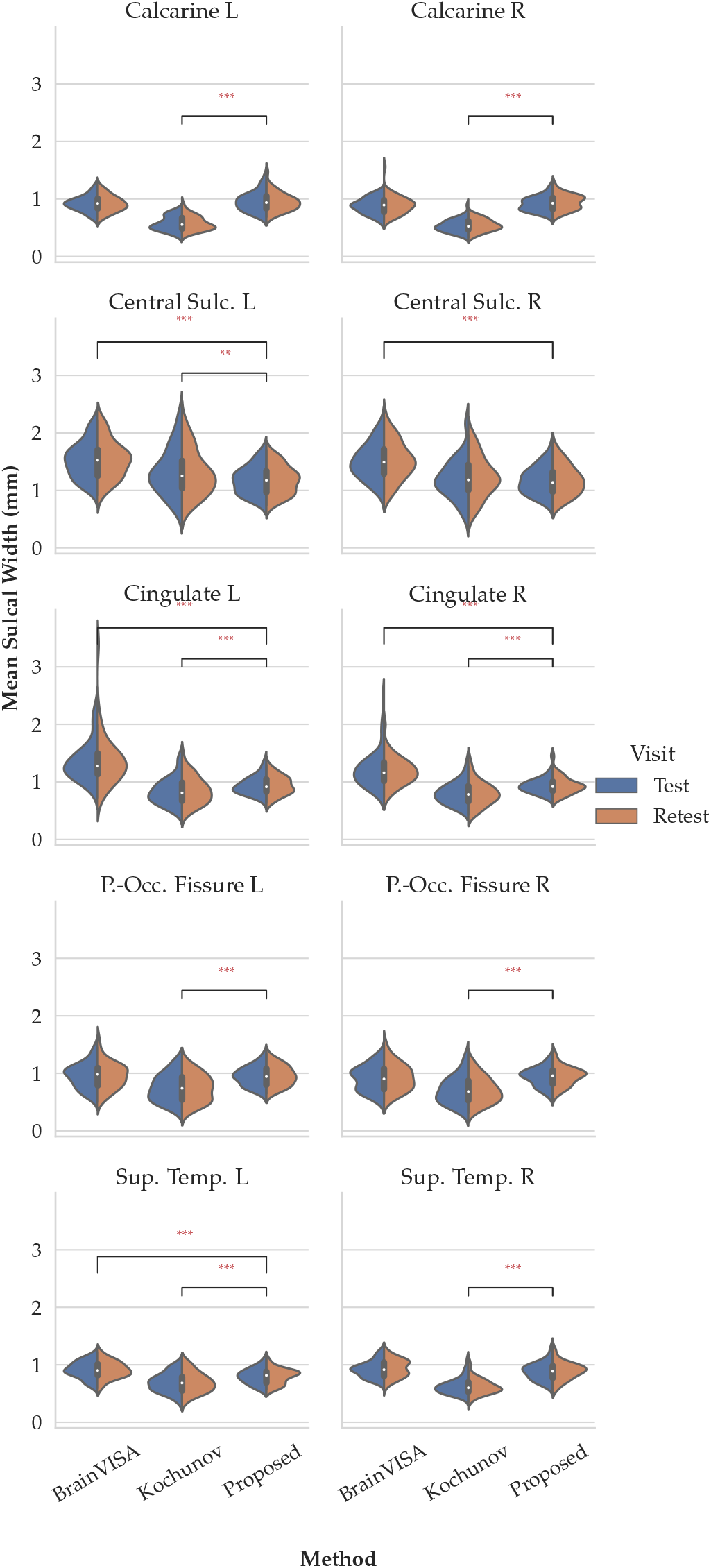
Violin plot representing the mean sulcal width for the ten selected sulci. For each method, the blue part of the plot represents the values obtained for the test acquisition, while the orange part represents the mean sulcal width values for the retest acquisition. Symbols “*”, “**”, and “***”, stand for significant differences between methods with FDR-corrected p-values lower than 0.05, 0.01, and 0.001 respectively.

**Table 1.** Sulcal width comparisons. Mean values and standard deviations are reported in millimeters. In bold the values that remained significant differences (*p <* 0.05) after FDR correction in the pairwise t-test.

| Sulcus Name | Proposed |  |  | Proposed - BrainVISA |  | Proposed - Kochunov |  |
| --- | --- | --- | --- | --- | --- | --- | --- |
|  | Mean (std) | BrainVISA Mean (std) | Kochunov Mean (std) | t (p-corrected) | Cohen's D (CI[95%]) | t (p-corrected) | Cohen's D (CI[95%]) |
| *Left Hemisphere* |  |  |  |  |  |  |  |
| Calcarine | 0.95 (0.16) | 0.92 (0.13) | 0.57 (0.12) | 2.04 (0.201) | 0.24 (-0.07, 0.54) | 19.85 (**<0.001**) | 2.72 (2.07, 3.38) |
| Central Sulcus | 1.18 (0.24) | 1.53 (0.32) | 1.31 (0.39) | -22.23 (**<0.001**) | -1.25 (-1.65, -0.85) | -3.47 (**0.006**) | -0.43 (-0.75, -0.12) |
| Cingulate | 0.94 (0.16) | 1.36 (0.39) | 0.84 (0.23) | -10.65 (**<0.001**) | -1.61 (-2.07, -1.16) | 4.83 (**<0.001**) | 0.50 (0.19, 0.82) |
| Parieto-Occipital Fissure | 0.95 (0.17) | 0.95 (0.22) | 0.74 (0.24) | -0.16 (1.000) | -0.02 (-0.32, 0.28) | 8.71 (**<0.001**) | 1.03 (0.65, 1.40) |
| Superior Temporal | 0.80 (0.13) | 0.91 (0.14) | 0.69 (0.17) | -7.35 (**<0.001**) | -0.83 (-1.18, -0.48) | 5.58 (**<0.001**) | 0.74 (0.40, 1.08) |
| *Right Hemisphere* |  |  |  |  |  |  |  |
| Calcarine | 0.93 (0.13) | 0.89 (0.15) | 0.54 (0.11) | 2.40 (0.098) | 0.31 (0.00, 0.61) | 26.64 (**<0.001**) | 3.45 (2.65, 4.24) |
| Central Sulcus | 1.18 (0.25) | 1.51 (0.31) | 1.22 (0.35) | -21.90 (**<0.001**) | -1.19 (-1.58, -0.80) | -1.62 (0.451) | -0.15 (-0.45, 0.16) |
| Cingulate | 0.94 (0.14) | 1.20 (0.29) | 0.80 (0.20) | -10.46 (**<0.001**) | -1.26 (-1.67, -0.86) | 9.52 (**<0.001**) | 0.81 (0.46, 1.15) |
| Parieto-Occipital Fissure | 0.94 (0.16) | 0.92 (0.22) | 0.71 (0.23) | 1.37 (0.676) | 0.12 (-0.18, 0.42) | 10.56 (**<0.001**) | 1.18 (0.79, 1.57) |
| Superior Temporal | 0.89 (0.15) | 0.93 (0.15) | 0.62 (0.14) | -2.09 (0.190) | -0.29 (-0.60, 0.01) | 12.80 (**<0.001**) | 1.89 (1.39, 2.39) |

Compared to the proposed method, BrainVISA displays higher values of width in the central sulcus (left: *p <* 0.001, *t* = − 22.23; right: *p <* 0.001, *t* = − 21.90) cingulate sulcus (left: *p <* 0.001, *t* = − 10.65; right: *p <* 0.001, *t* = − 10.46), and left superior temporal sulcus (*p <* 0.001, *t* = − 7.35), while the rest of comparisons are not significant.

The proposed method shows width values higher than the ones reported by Kochunov for the calcarine (left: *p <* 0.001, *t* = 19.85; right: *p <* 0.001, *t* = 26.64), cingulate (left: *p <* 0.001, *t* = 4.83; right: *p <* 0.001, *t* = 9.52), parieto-occipital fissure (left: *p <* 0.001, *t* = 8.71; right: *p <* 0.001, *t* = 10.56), and superior temporal sulcus (left: *p <* 0.001, *t* = 5.58; right: *p <* 0.001, *t* = 12.80). Kochunov reports higher values than the proposed method for the left central sulcus (*p* = 0.006, *t* = − 3.47).

The regressions between the proposed method and the other two methodologies (BrainVISA and Kochunov), along with Pearson’s correlation coefficient, are shown in Figure 5. On one hand, the correlation coefficient between BrainVISA and the proposed method is minimum in the left cingulate sulcus (*r* = 0.58, *p <* 0.001), while for left and right central sulcus, the correlation is *r* = 0.96 (*p <* 0.001). For the left and right parieto-occipital fissures *r* values are 0.80 and 0.84, respectively (both *p <* 0.01). On the other hand, the correlation coefficients between the proposed method and Kochunov are higher for the cingulate sulcus (left: *r* = 0.80, *p <* 0.001, right: *r* = 0.88, *p <* 0.001). Lowest correlation values between the proposed method and Kochunov are found in the superior temporal sulcus left: *r* = 0.61, *p <* 0.001; right: *r* = 0.52, *p <* 0.001.

**Fig. 5.**
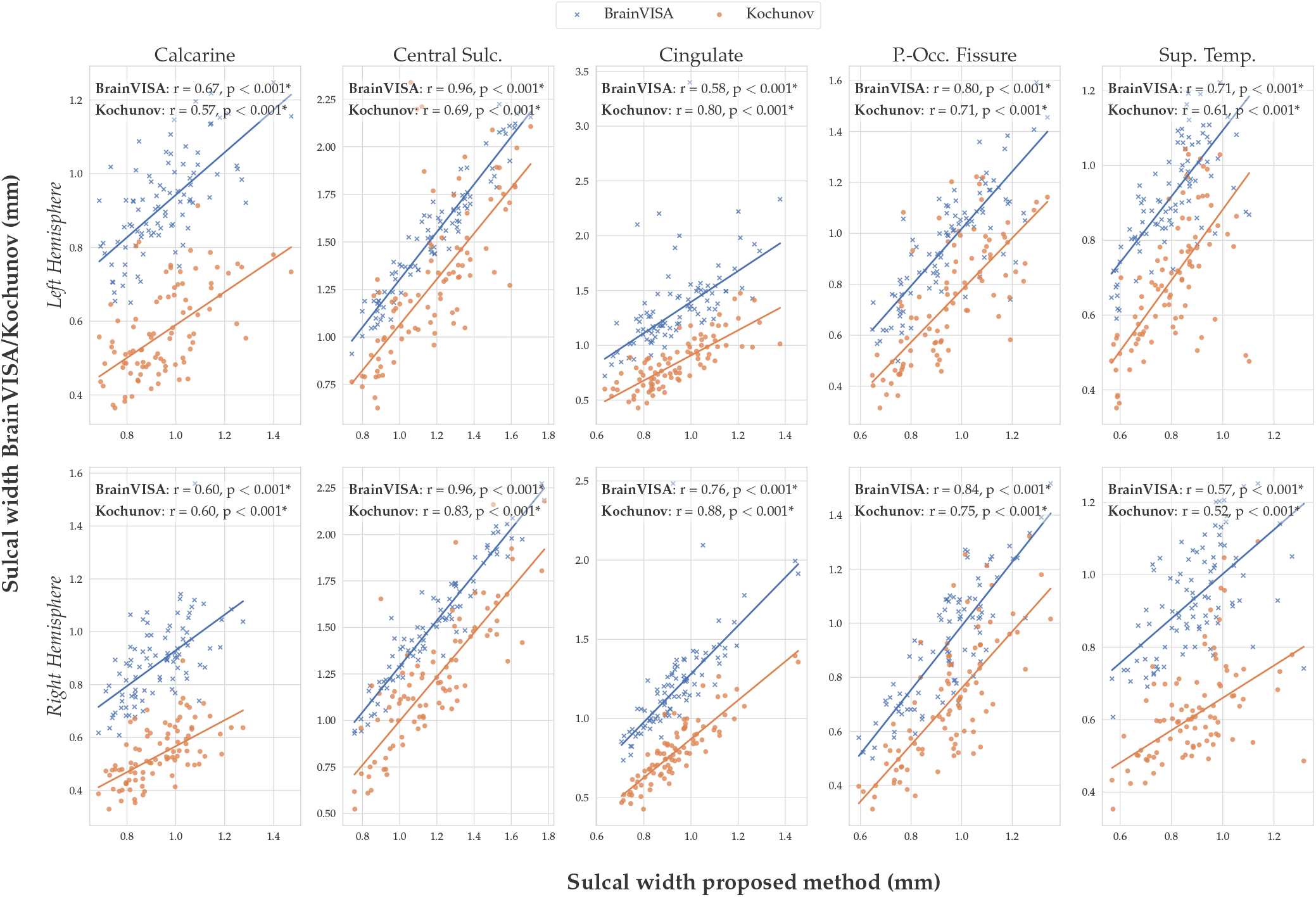
Width regression per sulcus. All p-values reported are corrected using FDR. Symbol “*” stands for significant values (*p <* 0.05) after correction. P.-Occ. Fissure stands for Parieto-Occipital Fissure. Sup. Temp. stands for Superior Temporal sulcus.

The ICC results (Table 2) indicate that the proposed method has a minimum ICC value for the right calcarine sulcus (ICC = 0.821) and maximum ICC value for the left central sulcus (ICC = 0.961). ICC for BrainVISA ranges between 0.484 (left cingulate sulcus) and 0.971 (left central sulcus). For Kochunov, the ICC values vary from 0.796 in the right calcarine sulcus to 0.946 in the left superior temporal sulcus. The results for every combined sulcus show that the proposed method’s ICC value is 0.930, while BrainVISA has an ICC of 0.869 and Kochunov scores 0.934.

**Table 2.** Intraclass correlation coefficient (ICC) estimating the resemblance between two different measures of a sulcus for the same subject. Bold values in p-corrected column indicate significant results (*p <* 0.05) after correcting for multiple comparisons using FDR.

| Sulcus Name | Proposed |  | BrainVISA |  | Kochunov |  |
| --- | --- | --- | --- | --- | --- | --- |
|  | ICC (CI[95%]) | p-corrected | ICC (CI[95%]) | p-corrected | ICC (CI[95%]) | p-corrected |
| *Left Hemisphere* |  |  |  |  |  |  |
| Calcarine | 0.862 (0.760, 0.920) | **<0.001** | 0.894 (0.820, 0.940) | **<0.001** | 0.854 (0.750, 0.920) | **<0.001** |
| Central Sulcus | 0.961 (0.930, 0.980) | **<0.001** | 0.971 (0.950, 0.980) | **<0.001** | 0.813 (0.690, 0.890) | **<0.001** |
| Cingulate | 0.886 (0.800, 0.940) | **<0.001** | 0.484 (0.220, 0.680) | **0.002** | 0.904 (0.830, 0.950) | **<0.001** |
| Parieto-Occipital Fissure | 0.896 (0.820, 0.940) | **<0.001** | 0.940 (0.890, 0.970) | **<0.001** | 0.944 (0.900, 0.970) | **<0.001** |
| Superior Temporal | 0.883 (0.800, 0.930) | **<0.001** | 0.877 (0.790, 0.930) | **<0.001** | 0.946 (0.900, 0.970) | **<0.001** |
| *Right Hemisphere* |  |  |  |  |  |  |
| Calcarine | 0.821 (0.660, 0.900) | **<0.001** | 0.678 (0.480, 0.810) | **<0.001** | 0.796 (0.660, 0.880) | **<0.001** |
| Central Sulcus | 0.954 (0.920, 0.970) | **<0.001** | 0.965 (0.940, 0.980) | **<0.001** | 0.845 (0.740, 0.910) | **<0.001** |
| Cingulate | 0.918 (0.850, 0.950) | **<0.001** | 0.710 (0.530, 0.830) | **<0.001** | 0.905 (0.830, 0.950) | **<0.001** |
| Parieto-Occipital Fissure | 0.890 (0.810, 0.940) | **<0.001** | 0.911 (0.840, 0.950) | **<0.001** | 0.942 (0.900, 0.970) | **<0.001** |
| Superior Temporal | 0.851 (0.740, 0.920) | **<0.001** | 0.837 (0.720, 0.910) | **<0.001** | 0.861 (0.760, 0.920) | **<0.001** |
| All | 0.930 (0.880, 0.960) | **<0.001** | 0.896 (0.820, 0.940) | **<0.001** | 0.934 (0.880, 0.960) | **<0.001** |

Finally, RMSE results (Table 3) indicate the proposed method scores lower errors for central sulcus (left: 0.005 mm; right: 0.006 mm), cingulate (left: 0.006 mm; right: 0.003 mm), and parieto-occipital fissure (left: 0.006 mm; right: 0.006 mm). BrainVISA shows lower errors for the left calcarine sulcus (0.004 mm), while Kochunov reports lower errors for the right calcarine sulcus (0.005 mm) and superior temporal sulcus (left: 0.003 mm, right: 0.005 mm). The combined value for every sulcus is lower for the proposed method with 0.006 mm.

**Table 3.** Root mean square errors (mm) between test and retest measures for each method and sulcus. Bold values indicate the minimum value for each row.

| Sulcus Name | Proposed | BrainVISA | Kochunov |
| --- | --- | --- | --- |
| *Left Hemisphere* |  |  |  |
| Calcarine | 0.007 | **0.004** | 0.004 |
| Central Sulcus | **0.005** | 0.006 | 0.058 |
| Cingulate | **0.006** | 0.156 | 0.010 |
| Parieto-Occipital Fissure | **0.006** | 0.006 | 0.006 |
| Superior Temporal | 0.004 | 0.005 | **0.003** |
| *Right Hemisphere* |  |  |  |
| Calcarine | 0.006 | 0.014 | **0.005** |
| Central Sulcus | **0.006** | 0.007 | 0.037 |
| Cingulate | **0.003** | 0.048 | 0.008 |
| Parieto-Occipital Fissure | **0.006** | 0.009 | 0.006 |
| Superior Temporal | 0.007 | 0.007 | **0.005** |
| All | **0.006** | 0.026 | 0.014 |

### Relationship Between Vertex-wise Sulcal Width and Age

Vertex-wise analyses revealed a significant positive correlation between age and sulcal width. Clusters surviving the multiple comparisons correction (CFT *<* 0.0001, CWP *<* 0.05) are those located in the central, superior temporal, and temporoparietal sulci, as well as the lateral fissure and some parts of the cingulate sulcus and the parieto-occipital fissure (see Figure 6).

**Fig. 6.**
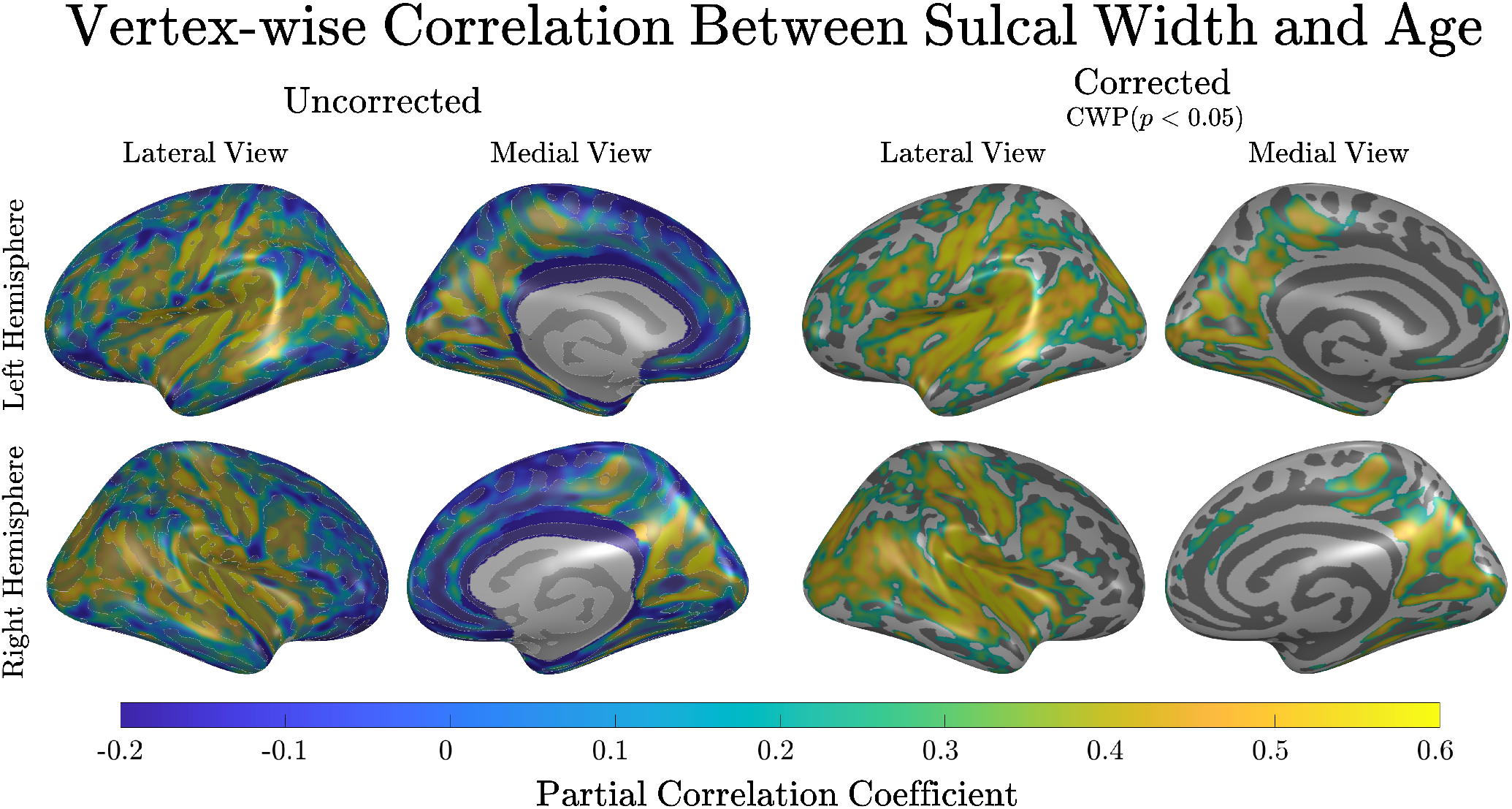
Vertex-wise partial correlation coefficients between age and width. The figures on the right side present results with a CFT of 0.0001 and a Cluster-Wise Probability (CWP) threshold of *p <* 0.05.

## Discussion

The presented results demonstrate that the proposed method provides a reliable vertex-wise measure of the physical distance between cortical sulcal banks. The algorithm, validated using simulated data and real data and compared against two widely-used sulcal width estimation algorithms, provides the community with an innovative method for creating cortical vertex-wise sulcal width maps.

First, validation using simulated data shows the accuracy of the results in a controlled environment. Secondly, using the HCP test-retest dataset and averaging vertex-wise values of sulcal width across ROIs, we enabled comparison with the ROI-based methods of BrainVISA (10) and Kochunov (13). Results from our algorithm displayed a high correlation with the values obtained using BrainVISA and Kochunov. In addition, the results indicate a robust reproducibility of our method, very similar to the one scored by the two mentioned toolboxes. Finally, the results from our group-wise vertex-wise analysis on OASIS-3 data extend prior findings from ROI studies by demonstrating individual, within-sulci variability in the positive association between age and sulcal width.

Validating sulcal width estimation methods is challenging as there is no available gold standard or ground truth. Therefore, by creating a simulated sulcus with known width for each vertex, we intended to approximate a ground truth to assess our method. Although the simulated sulcus does not reach the complexity of the most intricate sulci, it can mimic the characteristics of the regular sections of the sulcal basins, providing a well-described setting to evaluate the algorithm’s performance. Across the grand majority of vertices, our method produces width values that show a high correlation with the ground truth values. This provides the certainty that the algorithm properly measures widths for a simple representation of a sulcus.

We further validated our method using test-retest real data. Given the absence of real-life ground truth vertex-wise sulcal width data, we compared results from our method against results from existent ROI-based sulcal width toolboxes as BrainVISA (10), and Kochunov (13). Notably, the width values from these three approaches are similar for most sulci. Cross-method comparisons show, for example, that correlation coefficients between our method and Kochunov are between 0.52 and 0.88 for every sulcus, which is considerably higher than previous cross-method comparisons not involving our algorithm as the EDT-based approach in Mateos et al., that reports *r >* 0.5 only for the central sulcus. This analysis demonstrates that the proposed method behaves similarly to the reference methods at the ROI level. Furthermore, ICC values show excellent reproducibility for all the algorithms analyzed, except the left cingulate for BrainVISA. Precisely, the lower ICC value for BrainVISA in the left cingulate sulcus could explain the lower correlation between our method and BrainVISA for that ROI.

Having established that our method produces, at the ROI level, highly similar results to the ones delivered by Kochunov and BrainVISA, we now must mention that these two methods, along with others such as calcSulc (14), present a fundamental issue: they are bounded to a given atlas and provide averaged or median values for a whole region. The proposed method generates a vertex-wise map that boosts the number of analyses available for this metric. For instance, one can perform analyses in native space using any atlas of interest. Of course, vertex-wise studies are also an option, allowing the user to develop advanced models, e.g., investigate the relationship between changes in cortical thickness and sulcal width.

Employing the OASIS-3 healthy aging dataset, we used our method to analyze the vertex-wise association between age and sulcal width across individuals. The results reveal considerable variability of association strength across sulcal regions. This clearly demonstrates an advantage of our algorithm over ROI-based methods. That is, our method allows for the detection of within-sulci variability, whereas ROI-based methods only produce width averages that obscure within-sulci variability. The clusters of vertices that survived the multiple comparisons correction are located primarily in the central sulcus, superior frontal sulcus, inferior temporal sulcus, and lateral sulcus or parieto-occipital fissure. These results extend the findings of Jin et al. and Kochunov et al. by showing that the widening of sulci during senescence is widespread but also varies considerably within the ROIs.

## Limitations

The developed algorithm relies on identifying the different sulcal banks that compose each isoline. This critical step relies on detecting the points of high curvature of the curve based on a threshold. This threshold, chosen after visual inspection of different sulci in the dataset, is high enough to capture the most pronounced ones without including unnecessary divisions. However, fine-tuning might be needed when applying the method to populations with different characteristics, such as Alzheimer’s disease or infants. After segmenting the isoline curves into different sulcal banks, the next step is to identify the matching points on the opposite wall. Using the Euclidean distance to find this closest point presents one inconvenience: it tends to match points close to the division between sulcal banks with points adjacent to the division point instead of the one across the sulcus. This problem is noticeable when analyzing the results of the simulated sulcus, where errors are located in the regions near the wall division. However, this should not significantly affect the overall results since the error is limited to minimal areas near the junction between walls.

Finally, the results of vertex-wise analyses involving gyral regions must be addressed carefully, as those regions might not fit well in the sulcal width definition. These regions might be excluded from the analyses masking the gyral areas, providing only results for the sulcal regions to simplify the interpretation of the results.

## Conclusions

In light of the presented results, the developed method produces a reliable direct measure of the sulcal width, providing vertex-wise maps while tackling the main problems found in the previous literature. The algorithm and its source code are publicly available in https://github.com/HGGM-LIM/SWiM.

## Acknowledgments

This work was supported by the project exAScale Program-Ing models for extreme Data procEssing (ASPIDE), that has received funding from the European Union’s Horizon 2020 research and innovation program under grant agreement No 801091. This work has received funding from “la Caixa” Foundation under the project code LCF/PR/HR19/52160001. Susanna Carmona is funded by Instituto de Salud Carlos III, co-funded by European Social Fund “Investing in your future” (Miguel Servet Type I research contract CP16/00096). The CNIC is supported by the Instituto de Salud Carlos III (ISCIII), the Ministerio de Ciencia e Innovación (MCIN) and the Pro CNIC Foundation, and is a Severo Ochoa Center of Excellence (SEV-2015-0505).

Data were provided in part by the Human Connectome Project, WU-Minn Consortium (Principal Investigators: David Van Essen and Kamil Ugurbil; 1U54MH091657) funded by the 16 NIH Institutes and Centers that support the NIH Blueprint for Neuroscience Research; and by the McDonnell Center for Systems Neuroscience at Washington University.

Data were provided in part by OASIS-3 (Principal Investigators: T. Benzinger, D. Marcus, J. Morris; NIH P50 AG00561, P30 NS09857781, P01 AG026276, P01 AG003991, R01 AG043434, UL1 TR000448, R01 EB009352. AV-45 doses were provided by Avid Radiopharmaceuticals, a wholly owned subsidiary of Eli Lilly).

## Footnotes

1 https://github.com/nipy/mindboggle

2 https://github.com/alecjacobson/gptoolbox

3 https://db.humanconnectome.org/

4 https://www.oasis-brains.org/

