## Supplementary Material for "Estimation of Vertex-wise Sulcal Width Maps on Cortical Surfaces"

### 1. Depth step size parameter selection

A correct value of depth step for the generation of isolines is essential, as it allows a proper sampling of the surface without unnecessary increments in the processing time. In this case, the simulated sulcus is used for the estimation of an optimal depth step value. This surface has a mean edge length of 0.5 mm, so we estimate width maps with depth steps values ranging from the edge length (0.5 mm) to 1/50 the edge length (0.01 mm). The differences between estimated values and simulated ones are computed. Figure 1 shows the variation of mean error and standard deviation for the different step sizes evaluated.

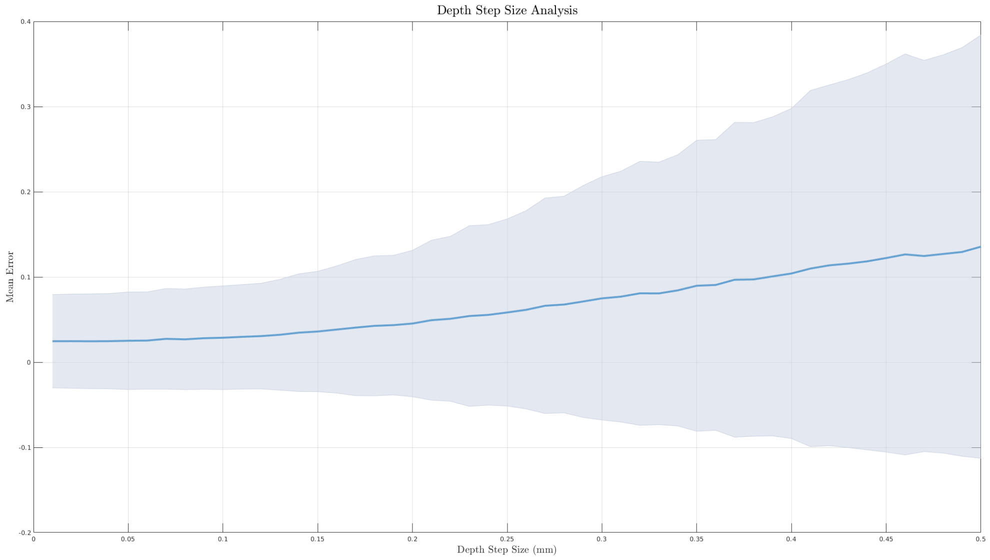


Figure 1: Plot of the mean error and its standard deviation for the different step sizes analyzed.

The results show a slope change or elbow approximately at 0.125 mm, which is 1/4 of the edge length. In this case, for the real data, we have edge lengths of 1 mm, meaning that the optimal step size would be 0.25 mm. Adding a small security margin, the default value selected for this method is 0.20 mm.

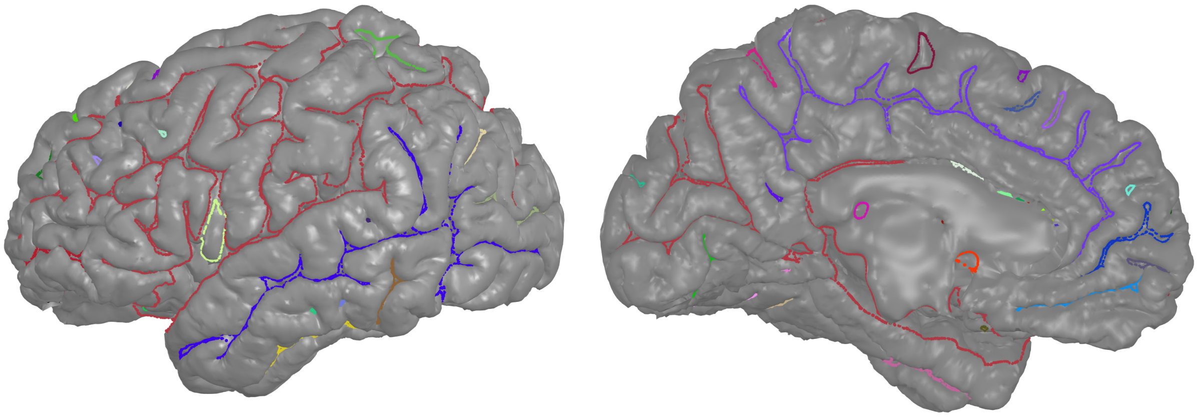

Figure 2: Frontal and medial view of a brain hemisphere depicting the isolines at a given depth level. Each color for the isolines represents a separate closed curve on the level.
